# An axon – T cell feedback loop enhances inflammation and axon degeneration

**DOI:** 10.1101/2023.01.06.523014

**Authors:** Tingting Liu, Huanhuan Wang, Daniel Y. Kutsovsky, Christine Y.J. Ohn, Nandan Patel, Jing Yang, David J. Simon

**Author notes:** equal contribution (Tingting Liu, Huanhuan Wang).

## Abstract

Inflammation is closely associated with many neurodegenerative disorders. Yet whether inflammation causes or exacerbates neurodegeneration has been challenging to define because the two processes are so closely linked. Here we disentangle inflammation from the axon damage it causes by individually blocking cytotoxic T cell function and axon degeneration. We model inflammatory damage in mouse skin, a barrier tissue that, despite frequent inflammation, must maintain proper functioning of a dense array of axon terminals. We show that sympathetic axons control skin inflammation through release of norepinephrine, which suppresses activation of gamma delta T cells via the β2 adrenergic receptor. Strong inflammatory stimulation in the form of the toll like receptor 7 (TLR7) agonist imiquimod (IMQ) causes progressive gamma delta T cell-mediated, Sarm-1-dependent loss of these immunosuppressive sympathetic axons, a positive feedback loop that removes a physiological brake on T cells, resulting in enhanced inflammation and inflammatory axon damage.

## Introduction

As a barrier tissue the skin serves two parallel and potentially conflicting roles. First is to enable precise sensation through the stereotyped placement of sensory axon terminals. Second is to protect the body from pathogens and physical injury. In this latter capacity skin is a site of frequent inflammation mediated by innate and adaptive immune responses (Pasparakis et al., 2014). Inflammation typically functions to protect tissues, however excessive inflammation can damage tissues, especially cells of the nervous system. Barrier axons are particularly vulnerable to inflammatory damage because they lack many of the key immune-protective mechanisms present in the central nervous system (Chiu et al., 2012; Croese et al., 2021; Ren and Dubner, 2010). Controlling the onset and magnitude of inflammation is therefore critical to barrier axon survival and function, and to neuronal survival in inflamed tissues more broadly.

Our understanding of the mechanisms that protect barrier axons from inflammatory damage and how these protections are overcome is limited. Indeed, whether inflammation initiates, amplifies, or is caused by neuronal loss has been the subject of extensive debate in multiple systems, in part because neuronal damage caused by inflammation obscures whether and how neurons contribute to inflammation (Martini and Willison, 2016). Are healthy neurons passively damaged by inflammatory mediators such as T cells, or do neurons actively recruit or repel these mediators? To disentangle these roles, we used a well-defined stimulus, the Toll-like receptor (TLR) 7 agonist imiquimod (IMQ), that activates a localized inflammatory response in mouse skin charactered by the appearance of interleukin-17 (IL-17)-producing γδ T cells. Skin-applied IMQ is widely used to model the inflammatory disorder psoriasis (Flutter and Nestle, 2013). In this context we observed progressive inflammation and axon loss in mouse dorsal skin. We used these hallmarks as a basis to test a panel of genetic and pharmacological manipulations that either block the cytotoxic function of activated T cells or that enhance or inhibit the resultant axon degeneration. Together these tools allowed us to study each aspect of T cell – axon signaling in isolation to define the signaling relationship between inflammation and axon loss in skin.

We show that inflammatory axon damage in mouse skin is controlled by a positive feedback loop between tyrosine hydroxylase (TH) positive sympathetic axons and cytotoxic γδ T cells. In healthy skin, sympathetic axons release norepinephrine which activates β2 adrenergic receptors (β2AR) on γδ T cells to inhibit their activation and expression of inflammatory cytokines. Yet once activated by IMQ, γδ T cells – and potentially additional cytotoxic lymphocytes – become cytotoxic towards sympathetic axons, driving their Sarm1-dependent degeneration, thus removing a key immunosuppressive axis. The net effect is an amplification of overall inflammation and an exacerbation of axon degeneration. Identification of this feed-forward inflammatory loop and the critical modulatory role of β2AR signaling has implications for the many neurodegenerative disorders accompanied by pronounced and progressive inflammation.

## Results

### Modeling inflammatory axon degeneration

Application of the Toll like receptor 7 agonist imiquimod (IMQ) to mouse skin is widely used as a method to cause inflammation and to model the inflammatory skin disorder psoriasis (Flutter and Nestle, 2013). We treated the dorsal skin of immunocompetent C57BL6/J mice with IMQ daily for six consecutive days (Figure 1A), which caused progressive inflammation, plaque-like psoriasis, and tissue pathology, as previously described (Figure S1A) (van der Fits et al., 2009). In this context we examined whether IMQ-dependent inflammation affected skin innervation. Here we developed a new skin preparation to visualize innervation over a large scale using whole mount immunolabeling and light sheet microscopy. We observed a dense reticulate structure of axons using the pan-neuronal marker PGP9.5 (Figure S1A), and a more ordered, brush-like distribution of tyrosine hydroxylase (TH) positive sympathetic axons (Figure 1B). Notably, treatment with IMQ caused a progressive decline in axon number, confirming our ability to induce inflammatory axon loss with spatial and temporal precision (Figure 1C).

**Figure 1.**
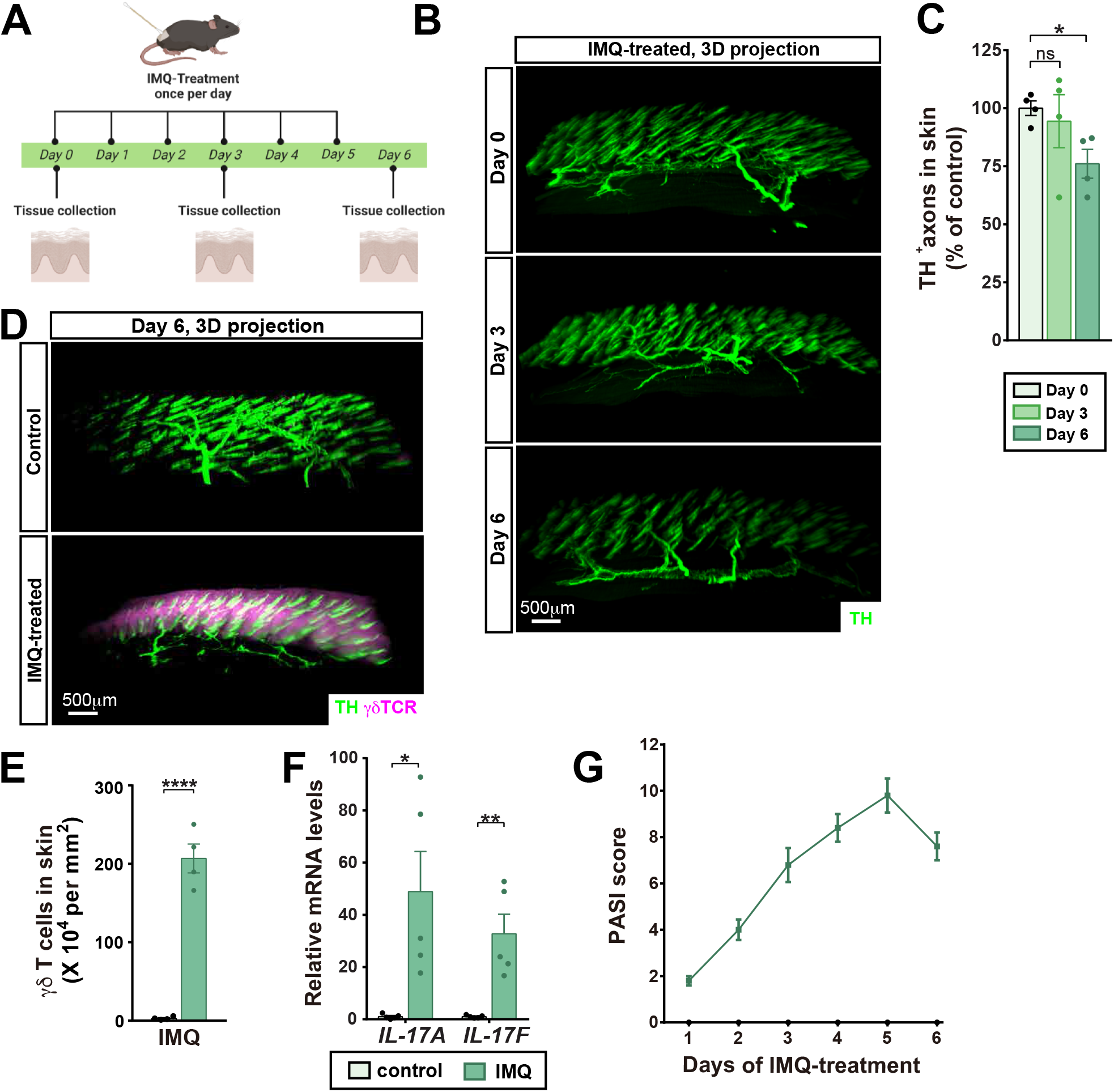
A model for T cell-mediated axon loss in mouse skin. (**A**) Mouse dorsal skin was treated with a non-toxic cream or 5% IMQ cream once per day for three or six consecutive days, with tissue collection as indicated. (**B**) Axonal projections throughout large sections of skin were visualized using light sheet microscopy followed by whole mount immunolabeling for TH. Representative 3D-projections of 1-mm thick skin samples are shown. (**C**) Axon density from (B) quantified for each time point as mean ± SEM. n = 4 animals. Here, and throughout: *p < 0.05, **p<0.01, ***p< 0.001, ****p< 0.0001 (unpaired, two-tailed t-test). (**D**) Recruitment of γδ T cells following IMQ treatment is visualized by whole mount immunolabeling. Representative 3Dprojections of 1 mm thick skin samples shown. (**E**) γδ-TCR-positive T cells were quantified from (D). n = 4 per condition, mean ± SEM, unpaired, two-tailed t-test. (**F**) Expression levels of *IL-17A* and *IL-17F* in skin samples collected 24 hours after the last IMQ-treatment were determined by the qPCR. *n* = 5, mean ± SEM, unpaired, two-tailed t-test. (**G**) PASI scoring of mouse dorsal skin was evaluated every day before the subsequent IMQ-treatment. *n* = 5 mice, means ± SEM.

To investigate the mechanism of IMQ-induced axon loss we focused on TH^+^ axons. We observed a large increase in γδ T lymphocytes in close proximity to TH^+^ axons (Figure 1D,E), a corresponding increase in mRNA abundance of their principal cytokines IL-17A and IL-17F (Figure1F), and progressive psoriasis-like skin pathology (Figure 1G). To test whether T lymphocytes drive the observed axon loss, we repeated IMQ treatment in *Tcrbd* mice which lack the a, β, and γδ T-cell receptors (Itohara et al., 1993; Mombaerts et al., 1991) and therefore are deficient in all T lymphocyte populations. We found that knockout mice had significantly less axon loss when challenged with IMQ (Figure 2A,B), consistent with a lack of T cell-mediated axon damage. Further consistent with a lack of damaging T cells (Figure 2F), these mice displayed diminished IMQ-induced tissue pathology (Figure 2C-E).

**Figure 2.**
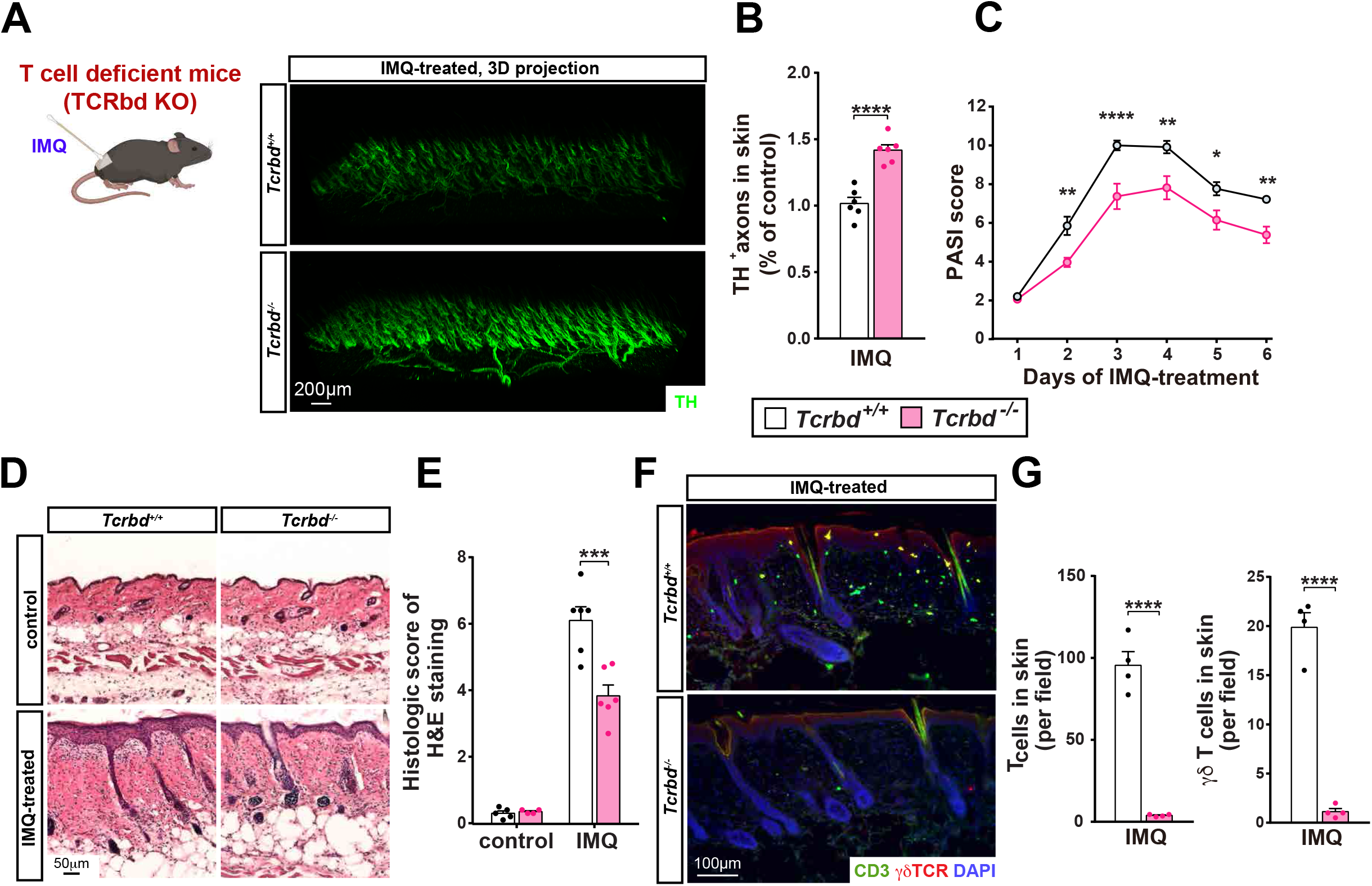
T cells are required for IMQ-mediated axon loss and tissue damage. (**A**) Mice of the indicated genotypes were subjected to six days of daily IMQ treatment. TH^+^ axons were visualized at the terminal time point, and their density was quantified (**B**). *n* = 6, mean ± SEM, unpaired, two-tailed t-test. The response to IMQ was further assayed by PASI (**C**) *n* = 6, mean ± SEM, two-way ANOVA with Sidak’s correction. (**D**) IMQ-induced tissue damage was assessed by H&E and quantified using Baker’s standard (**E**). n = 4 (control) or 6 (IMQ), mean ± SEM, twoway ANOVA with Tukey’s post-hoc test. (**F**) Inflammation was assessed in tissue sections of the indicated genotypes and qualified (**G**) n = 4, mean ± SEM, unpaired, two-tailed t-test.

A prominent mode of T lymphocyte toxicity involves formation of cytotoxic pores in target cell plasma membranes through which granzymes and other cytotoxic molecules enter and trigger target cell death (Russell and Ley, 2002). Perforins are the major pore forming proteins found in cytotoxic T lymphocytes (CTLs), γδ T cells, and natural killer cells (Voskoboinik et al., 2015). Notably, deletion of perforin1 in *Prf1* knockout mice (Kagi et al., 1994) mitigated IMQ-mediated axon loss (Figure 3A,B), psoriasis (Figure 3C), and skin pathology (Figure 3D,E). Together these results indicate that IMQ-mediated TH^+^ axon loss is controlled by Perforin-1-mediated cytotoxicity by T lymphocytes, likely though perhaps not exclusively of the γδ T cell class.

**Figure 3.**
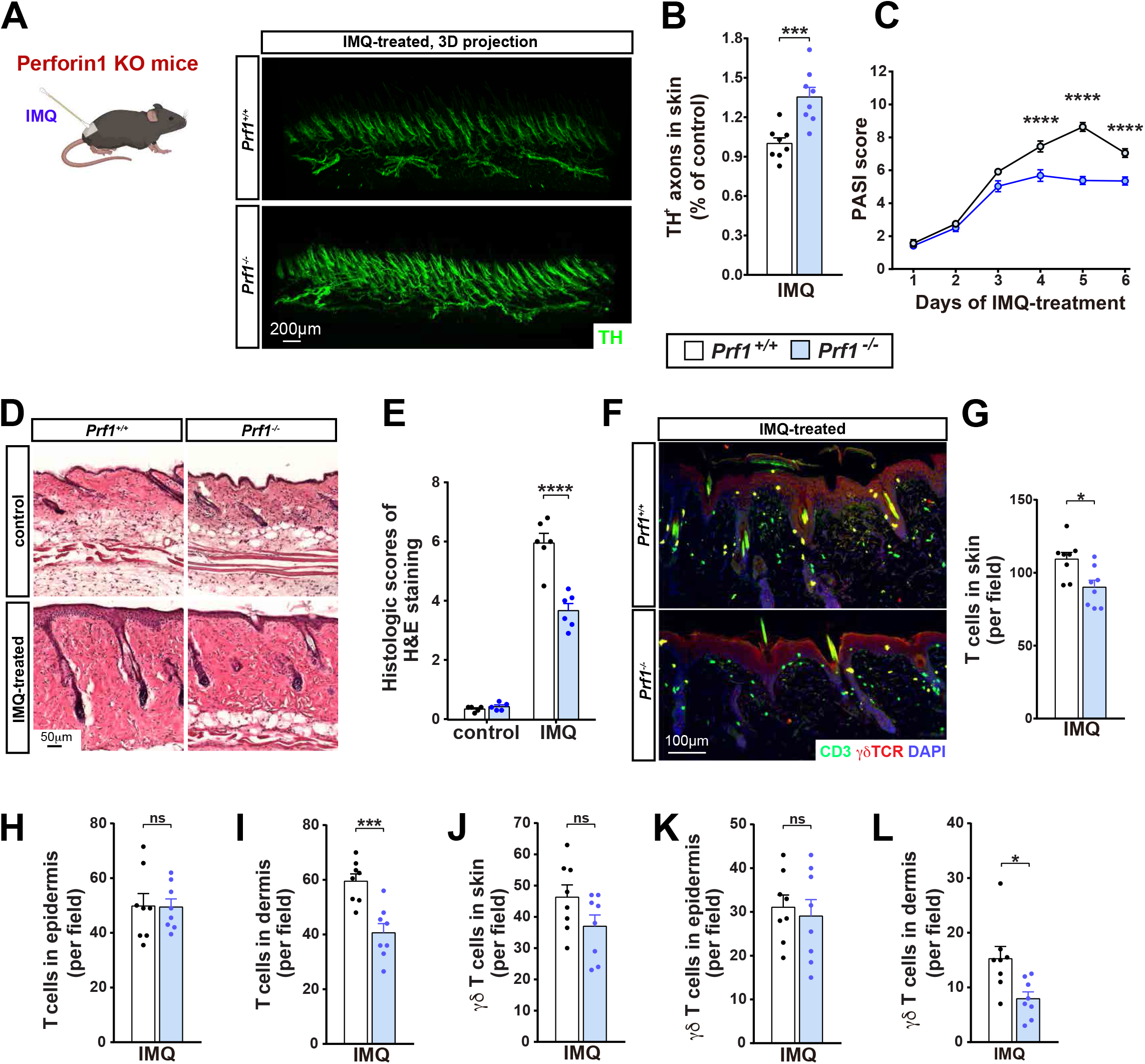
Perforin-1 is required for IMQ-mediated axon loss and tissue damage. (**A**) Mice of the indicated genotypes were subjected to six days of daily IMQ treatment. TH^+^ axons were visualized at the terminal time point, and their density was quantified (**B**). n = 8 mice, mean ± SEM, unpaired, two-tailed t-test. The response to IMQ was further assayed by PASI (**C**) *n* = 6, mean ± SEM, two-way ANOVA with Sidak’s correction. (**D**) IMQ-induced tissue damage was assessed by H&E and quantified using Baker’s standard (**E**). n = 5 (control) or 6 (IMQ), mean ± SEM, twoway ANOVA with Tukey’s post-hoc test. (**F**) Inflammation was assessed by immunostaining in tissue sections of the indicated genotypes. (**G-L**) The indicated populations were counted across each field of view at 10x magnification. Dermis and epidermis were identified by the density of the DAPI signal (F, blue channel) and differentially quantified. n = 8 mice, mean ± SEM, unpaired, two-tailed t-test.

Surprisingly, overall T cell number was lower in IMQ treated Perforin-1 knockouts (Figure 3F,G), contrary to our expectation that the protective phenotype of these knockouts would exclusively relate to cytotoxicity, not the degree of inflammation. Moreover, the reduction of both total T cells and the γδ T cell subset was specific to the dermal skin layer (Figure 3H-L), consistent with exclusive the projection of TH^+^ axons to the dermis, not the epidermis (Roosterman et al., 2006). These data drove us to consider whether IMQ-mediated axon loss might itself affect the inflammatory response. To test this hypothesis, we depleted TH^+^ axons in skin through direct application of the neurotoxic dopamine derivative 6-hydroxydopamine (6OHDA) and confirmed their elimination via immunostaining (Figure 4A,B). We observed that pre-treatment with 6OHDA enhanced IMQ-mediated γδ T lymphocyte influx (Figure 4G), expression of IL-17A and F (Figure 4C), and tissue pathology (Figure 4D-F). Importantly, 6OHDA treatment in the absence of IMQ did not affect any of these parameters.

**Figure 4.**
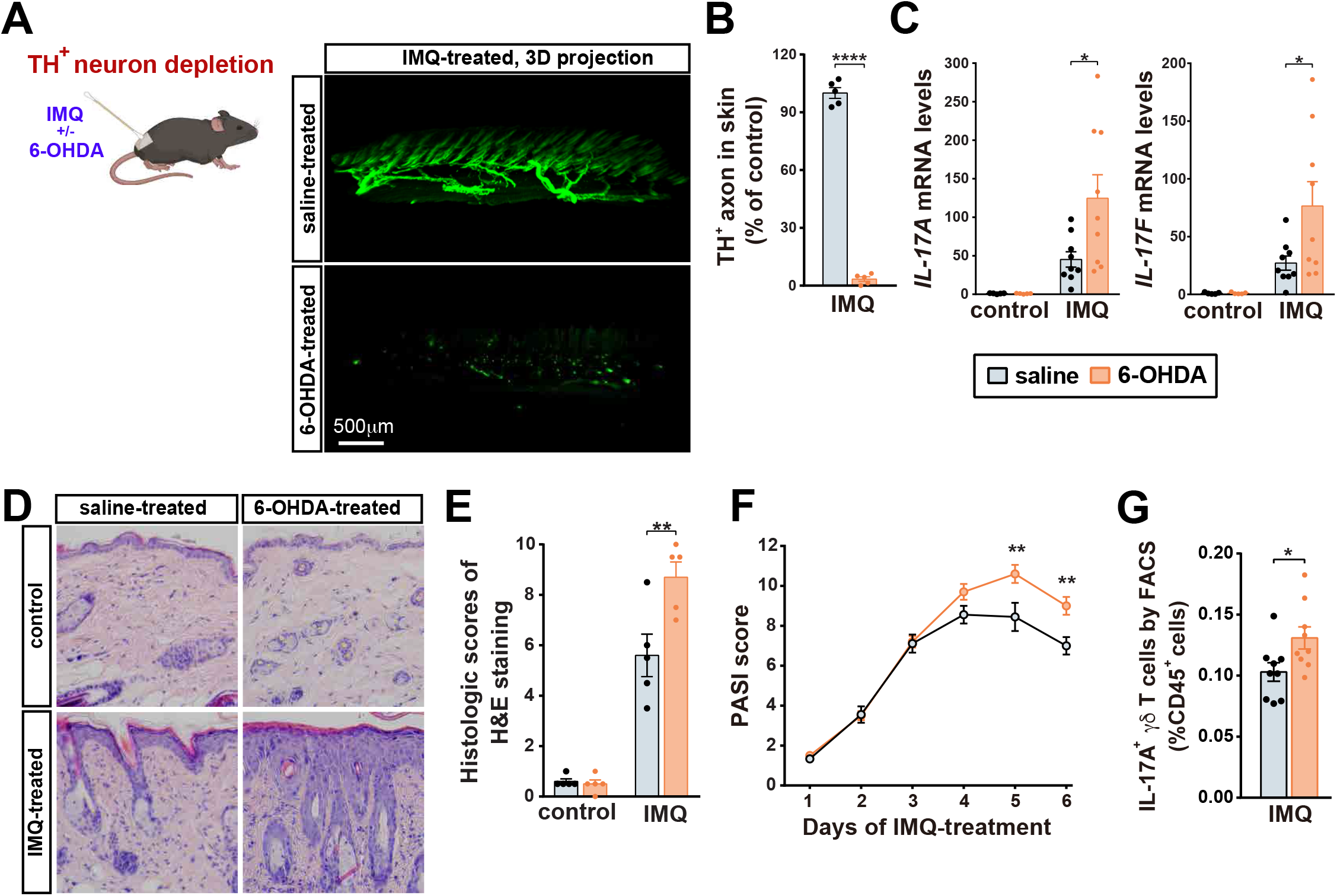
TH^+^ axon depletion enhances IMQ-induced skin inflammation. (**A**) Representative 3D-projection of skin treated as indicated and visualized by whole mount immunolabeling for TH. (**B**) TH-positive sympathetic axons were quantified. n = 5 mice, mean ± SEM, unpaired, two-tailed t-test. (**C**) Expression of *IL-17A* and *IL-17F* in skin was quantified by qPCR. *n* = 5 (control) or 9 (IMQ), mean ± SEM, two-way ANOVA with Tukey’s post-hoc test. (**D**) Representative images of the H&E-stained skin sections from the indicated conditions, quantified in (**E**) n = 5, mean ± SEM, two-way ANOVA with Tukey’s post-hoc test. (**F**) PASI of mouse dorsal skin was evaluated every day before IMQ-treatment. *n* = 9 mice, mean ± SEM, two-way ANOVA with Sidak’s correction. (**G**) CD45+ CD3+ γδTCR+ IL-17A+ cells in the skin tissue were quantified by flow cytometry. *n* = 9 mice, mean ± SEM, unpaired, two-tailed t-test.

That depletion of TH^+^ axons enhanced IMQ-mediated inflammation suggested that these axons are immunosuppressive in dorsal skin. To test this, we sought to measure inflammation in the absence of the axon damage that it normally causes. Sarm1 is an NAD^+^ hydrolyzing enzyme (Essuman et al., 2017) that governs a pro-degenerative pathway in axons (Sambashivan and Freeman, 2021). Its loss protects axons from degeneration in many contexts, including downstream of inflammatory cytokines (Ko et al., 2020; Sun et al., 2021). We found that IMQ-mediated axon loss was decreased in *Sarm1* knockout mice, and that this corresponded to a significant reduction in γδ T cell number and IL-17A and F expression in skin, as well as diminished psoriasis-like tissue pathology (Figure S2). While the precise mechanism linking Perforin-1-mediated rupture of the axonal plasma membrane to activation of Sarm1-mediated axon degeneration is unknown, our results nevertheless suggest that *Sarm1* knockout is a powerful tool to block IMQ-mediated axon loss.

Sarm1 is highly expressed in neurons, however its expression has been noted in other tissues, including immune cells (Jin et al., 2022; Panneerselvam et al., 2013). To specifically delete Sarm1 in TH^+^ neurons we crossed a Sarm1 conditional knockout allele to TH-Cre (*Th-Cre; Sarm1^fl/fl^*) mice. Similar to germline *Sarm1* knockout mice, TH^+^ axon density was higher following IMQ treatment in *Th-Cre; Sarm1^fl/fl^* skin (Figure 5A,B). Neuronal deletion of Sarm1 similarly lessened the degree of psoriasis (Figure 5C), tissue pathology (Figure 5D,E), and IMQ-induced IL-17A and F induction (Figure 5F). Notably, protection against psoriasis-like pathology was greater in these mice compared to germline *Sarm1^-/-^* mice (Figure S2H). Together these data demonstrate that TH^+^ axons are immunosuppressive and that their degeneration enhances skin ongoing inflammation.

**Figure 5.**
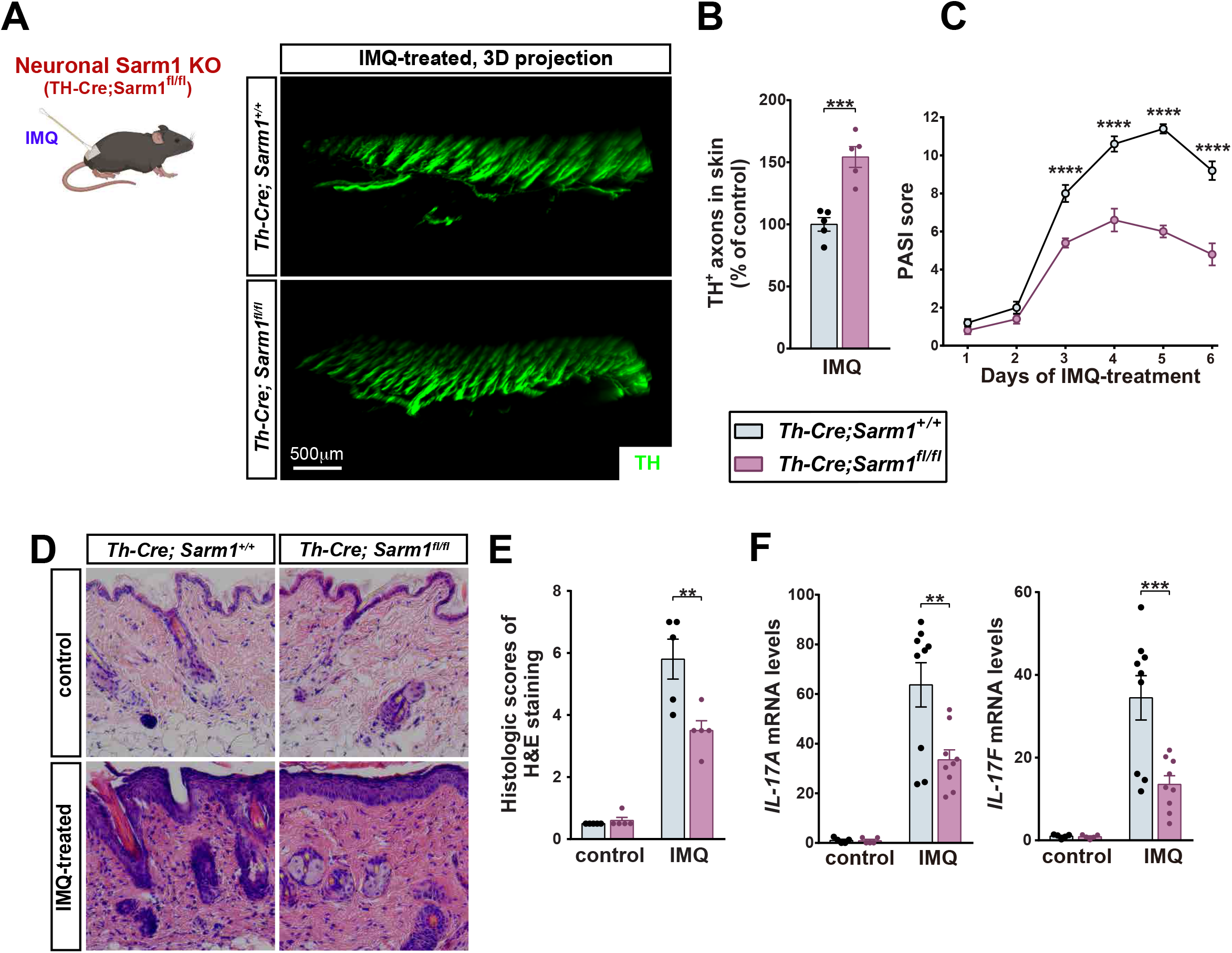
Blocking TH^+^ axon degeneration lessens IMQ-induced skin inflammation. (**A**) Mice of the indicated genotypes were subjected to six days of daily IMQ treatment. TH^+^ axons were visualized at the terminal time point, and their density was quantified (**B**). n = 5 mice, mean ± SEM, unpaired, two-tailed t-test. The response to IMQ was further assayed by PASI (**C**) *n* = 5 mice, mean ± SEM, two-way ANOVA with Sidak’s correction. (**D**) IMQ-induced tissue damage was assessed by H&E and quantified using Baker’s standard (**E**). n = 5 mice, mean ± SEM, twoway ANOVA with Tukey’s post-hoc test (**F**) Expression of *IL-17A* and *IL-17F* in skin was quantified by qPCR. *n* = 5 (control) or 9 (IMQ), mean ± SEM, two-way ANOVA with Tukey’s post-hoc test.

How do TH^+^ axons normally suppress skin inflammation? Norepinephrine (NE) is the main neurotransmitter released by TH^+^ sympathetic nerve terminals, and can have both immunosuppressive and immunostimulatory roles, depending on the cellular context (Gaskill and Khoshbouei, 2022). Primary γδ T cells were purified by flow cytometry from subcutaneous lymph nodes after IMQ dosing and treated *in vitro* with NE, which inhibited expression of activation markers IL-17A and F (Figure 6A). Subsequent quantification of mRNA expression of all adrenergic receptors in these acutely isolated γδ T cells identified the β2-adrenergic (Adrb2) receptor as most abundantly expressed (Figure 6B). Consistent with this expression pattern, treatment of primary γδ T cells with β2-adrenergic receptor agonists formoterol or clenbuterol each down-regulated IL-17A and F expression (Figure 6C,D). These data are consistent with axon-derived NE signaling via β2 adrenergic receptors on γδ T cells to suppress inflammation in dorsal skin.

**Figure 6.**
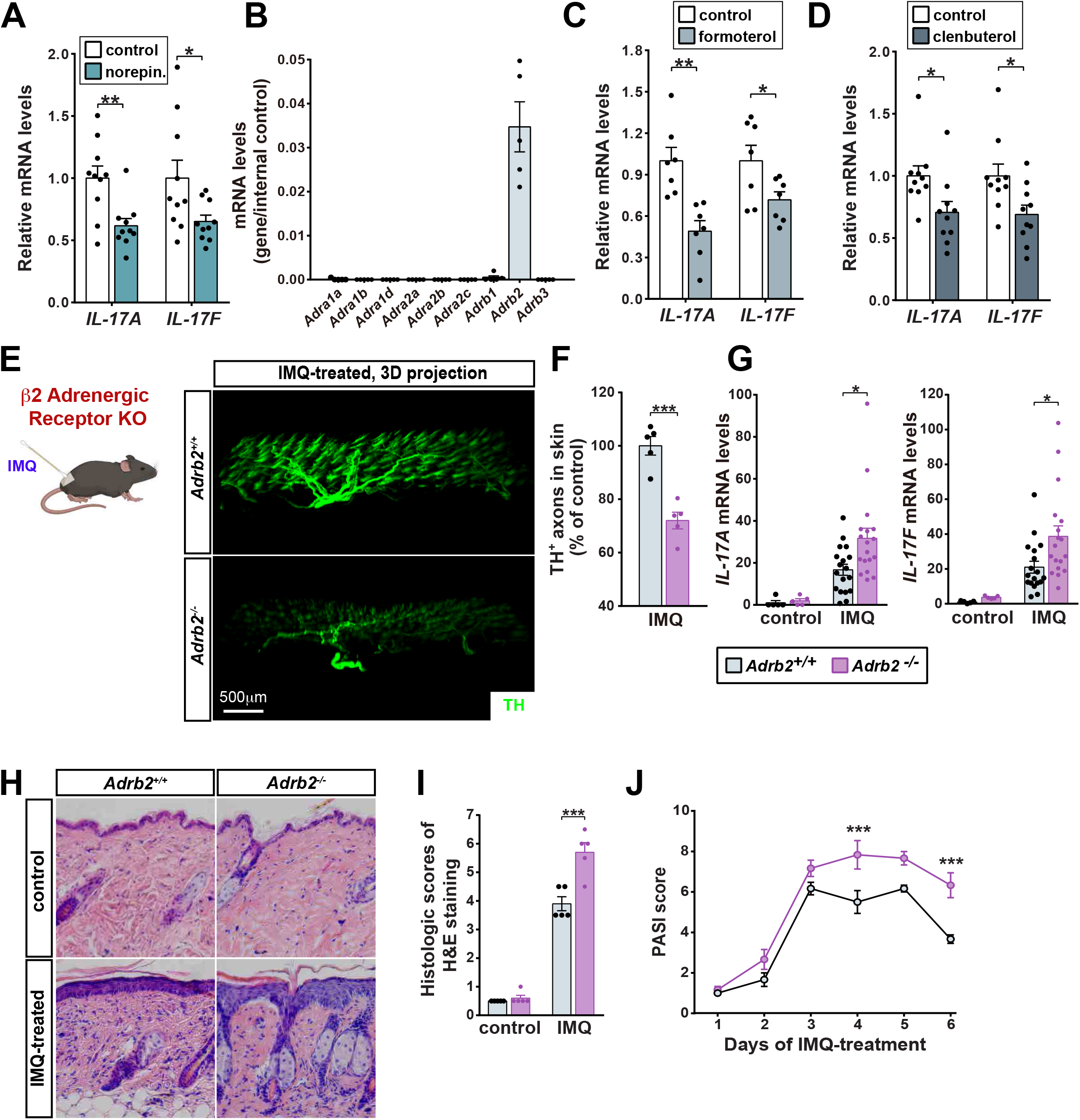
A norepinephring-β2-adrenergic receptor signaling axis lowers skin inflammation. (**A)** γδ T cells (CD45+ CD3+ γδTCR+) isolated from subcutaneous lymph nodes of IMQ-treated wild-type mice were treated with 20 μM NE, followed by expression analysis of indicated genes. *n* = 10 mice, mean ± SEM, unpaired, two-tailed t-test (**B**) The expression profile of adrenergic receptors in γδ T cells was determined by qPCR. n = 5 mice, mean ± SEM. (**C, D**) γδ T cells were treated with 20 μM of the indicated drugs followed by expression analysis. *n* = 7 (formoterol) or 10 (clenbuterol), mean ± SEM, unpaired, two-tailed t-test (**E**) Representative 3D-projections images of 1-mm thick IMQ-treated skin sections of the indicated genotypes. TH^+^ axon density is quantified in (**F**). n = 5 mice, mean ± SEM, unpaired, two-tailed t-test. (**G**) Expression of *IL-17A* and *IL-17F* in skin was determined by qPCR. *n* = 5 (control) or 18 (IMQ), mean ± SEM two-way ANOVA with Tukey’s post-hoc test. (**H**) IMQ-induced tissue damage was assessed by H&E and quantified using Baker’s standard (**I**). n = 5 mice, mean ± SEM, two-way ANOVA with Tukey’s post-hoc test. (**J**) The response to IMQ was further assayed by PASI. *n* = 6 mice, mean ± SEM, two-way ANOVA with Sidak’s correction.

That β2-adrenergic receptor agonism dampened γδ T cell activation suggested that inhibition of this immunosuppressive pathway would heighten inflammation. Indeed, a major clinical side effect of β2-adrenergic receptor antagonists is psoriasis (Gold et al., 1988). We therefore examined skin immune responses induced by IMQ in germline *Adrb2* knockout mice. Notably, loss of β2-adrenergic receptor signaling significantly exacerbated TH^+^ axon loss (Figure 6E,F), expression of IL-17A and F (Figure 6G), as well as overall inflammation and psoriasis-like pathology (Figure 6H-J), similar to TH^+^ axon depletion using 6-OHDA treatment (Figure 4).

To determine the locus of β2-adrenergic receptor action we generated bone-marrow chimeric mice (BMCMs) by transplanting bone marrow of *Adrb2* wild type or knockout mice into wild type mice whose own bone marrow had been eliminated by irradiation (Figure 7A). Mice lacking Adrb2 in the bone marrow compartment (where γδ T cells originate) phenocopied germline *Adrb2* knockout mice with respect to axon loss (Figure 7B,C), IL-17A and F induction (Figure 7D), and psoriasis-like skin pathology (Figure 7E,G). While the bone marrow niche produces immune cells in addition to γδ T cells, this result nevertheless suggests that Adrb2 functions in the immune system rather than in neurons. Together our data support the existence of a feed-forward neuro-immune loop in mouse dorsal skin. Normally, NE released by TH^+^ axons inhibits skin inflammation via the β2-adrenergic receptor on γδ T cells. Cytotoxic damage to these same axons drives their Sarm1-dependent degeneration and ultimately diminishes their immunosuppressive capacity, to enhance inflammation and axon degeneration (Figure 7H).

**Figure 7.**
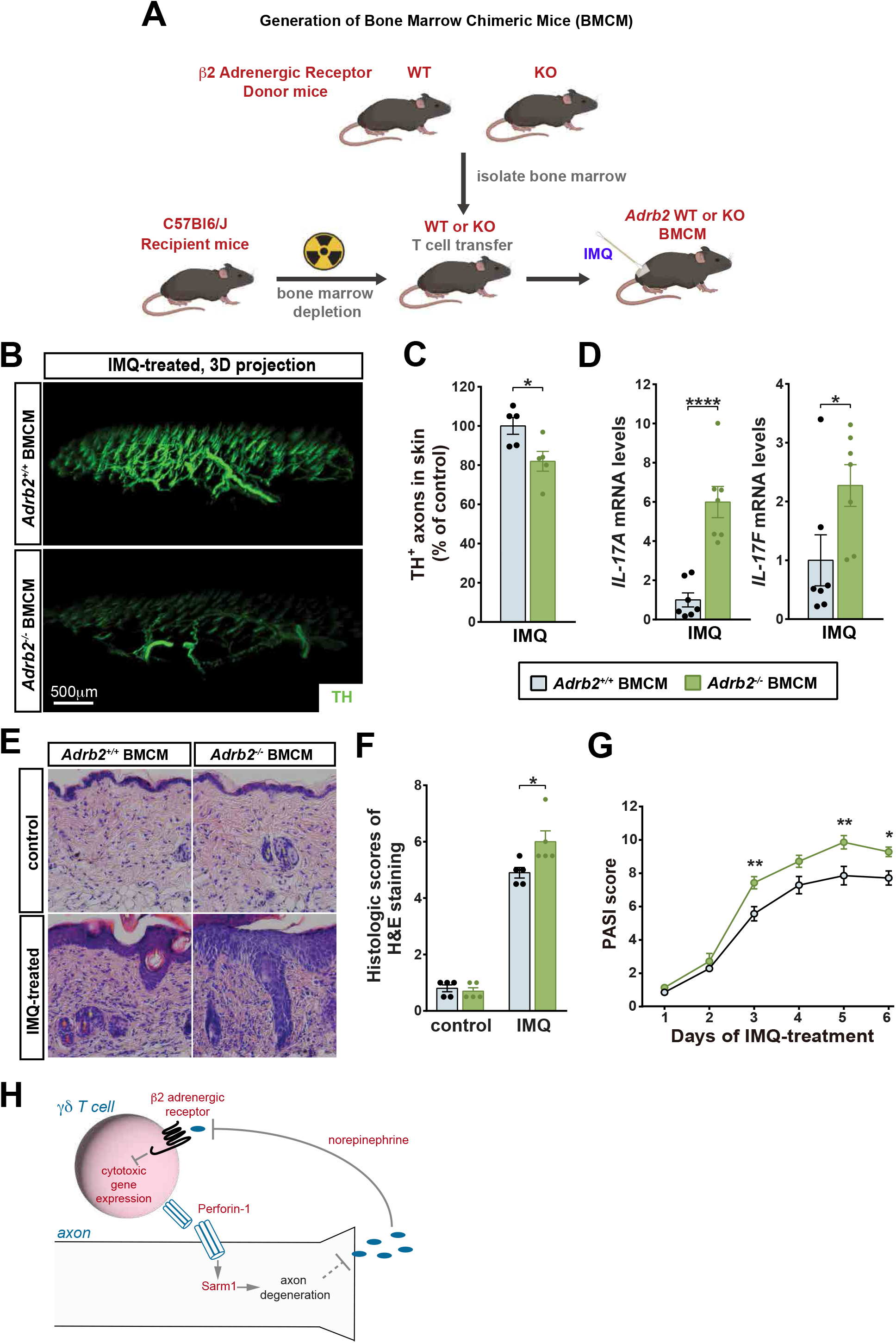
β2-adrenergic receptor acts in the immune compartment to limit IMQ-induced inflammation. (**A**) Schematic of bone marrow transplantation to generate Adrb2 chimeric mice. Wild type recipient mice were irradiated to deplete endogenous bone marrow, and subsequently injected with 2 × 10^6^ bone-marrow-derived cells from donor mice (*Adrb2^+/+^* or *Adrb2^-/-^*) via intravenous injection. Bone marrow chimeric mice (BMCM) were treated with IMQ (**B**) and TH^+^ axon number was quantified (**C**). n = 5 mice, mean ± SEM, unpaired, two-tailed t-test. (**D**) Expression level of *IL-17A* and *IL-17F* in skin was determined by the qPCR. *n* = 7 mice, mean ± SEM unpaired, two-tailed t-test. (**E**) Skin samples from the indicated conditions were processed for H&E staining and histologic scores were determined using Baker’s standard (**F**). n = 5, mean ± SEM, two-way ANOVA with Tukey’s post-hoc test. (**G**) The response to IMQ was further assayed by PASI. *n* = 7 mice, mean ± SEM, two-way ANOVA with Sidak’s correction. **(H)** We identify an axon-T-cell positive feedback loop that enhances inflammation and inflammatory axon damage.

### Discussion

Precise control of immune responses is critical to the proper function of barrier tissues and to nervous system function more broadly. Here we focused on TH^+^ axons in skin, however inflammation occurs throughout the nervous system and accompanies many neurodegenerative disorders and traumas that target diverse neuronal and non-neuronal cell types. There are many triggers of inflammation, both physiological and foreign, involving extensive interplay between multiple cells. Together these interactions complicate our ability to model inflammatory damage in the nervous system and to dissects its underlying mechanisms.

We use a simplified system to trigger acute inflammation and cause axon damage. IMQ is well characterized to activate resident IL-17-expressing γδ T cells in skin, resulting in tissue damage that is used as a pre-clinical model of psoriasis (Flutter and Nestle, 2013). We reasoned that application of IMQ would enable precise control over the timing, location, and cellular composition of inflammation, and allow a controlled setting to study inflammatory axon degeneration. Indeed, we find that application of IMQ to mouse dorsal skin causes progressive, T cell-dependent axon degeneration. In the context of this model, we identify a pro-degenerative positive feedback loop that reinforces inflammatory damage. Blocking IMQ-mediated axon damage by either impairing the cytotoxic function of activated T cells or by removing an axon-intrinsic pro-degenerative pathway in TH^+^ axons both lessen overall inflammation. Further, forcible ablation of TH^+^ axons (Figure 5) enhances IMQ-mediated skin inflammation. Together these data argue that TH^+^ axons normally suppress skin inflammation and that their loss enhances inflammation.

Dying cells release immune-stimulatory molecules that promote the clearance of cellular debris and the maintenance of tissue homeostasis (Yatim et al., 2017). While we do not rule out the importance of these signals in exacerbating IMQ-mediated axon loss, our data argue that disinhibition of an immunosuppressive β2 adrenergic receptor signaling axis, resulting from degeneration of TH^+^ axons, underlies the inflammatory axon damage that we describe. We show that norepinephrine, a neuromodulator known to be released from TH^+^ axons, suppresses γδ T cell activation. Further, deletion of its receptor, β2AR, in the immune compartment phenocopies forcible TH^+^ axon deletion and enhances IMQ-mediated inflammation. These data are consistent with a NE-β2AR signaling axis as a physiological brake on skin inflammation that, when interrupted, enhances inflammation and subsequent inflammatory axon damage. While our data support a prominent role for γδ T cells, other Perforin-1-expressing cytotoxic lymphocyte populations, such as CD8 T cells, may also contribute to the inflammatory feedback loop that we observe.

Positive feedback loops such as the one we define can predispose towards rapid and uncontrolled amplification (Ingolia and Murray, 2007), which may account for the progressive inflammation and neuronal loss that characterize many neurodegenerative pathologies. Control over the initiation and early amplification of inflammation would therefore be critical determinants of whether these loops are engaged. Intriguingly, sensory neurons themselves can drive early immune responses through the release of pro-inflammatory chemokines (Wang et al., 2018; Wlaschin et al., 2018) and other immune-stimulating peptides (Klein Wolterink et al., 2022), thus providing additional context to regulate the dynamics of resultant inflammatory loops. Finally, our findings highlight the importance of counteracting positive feedback loops to resolve inflammatory responses, especially when endogenous immunosuppressive mechanisms such as regulatory T cells have been exhausted. In skin, β2 adrenergic receptor agonism would likely dampen inflammatory responses by counteracting the loss of sympathetic axons. Elsewhere, direct engagement of inhibitory receptors on infiltrating lymphocytes or on activated CNS-resident immune cells may serve a similar role to counteract neuroinflammation in the context of neurodegenerative disorders.

## Acknowledgements

This work was supported by startup funds from Weill Cornell Medicine (D.J.S) and by the Fernholz Family Foundation (D.J.S). We thank members of the Simon lab for helpful comments.

## Author Contributions

Project conception and experimental design: T.L, D.J.S.; Experimentation: T.L, H.W, D.K, C.Y.J.O, N.P; Data analysis: T.L, D.J.S; Supervision: D.J.S, J.Y. Writing: T.L, D.J.S.

## Competing interests

None

---

**Figure S1.**
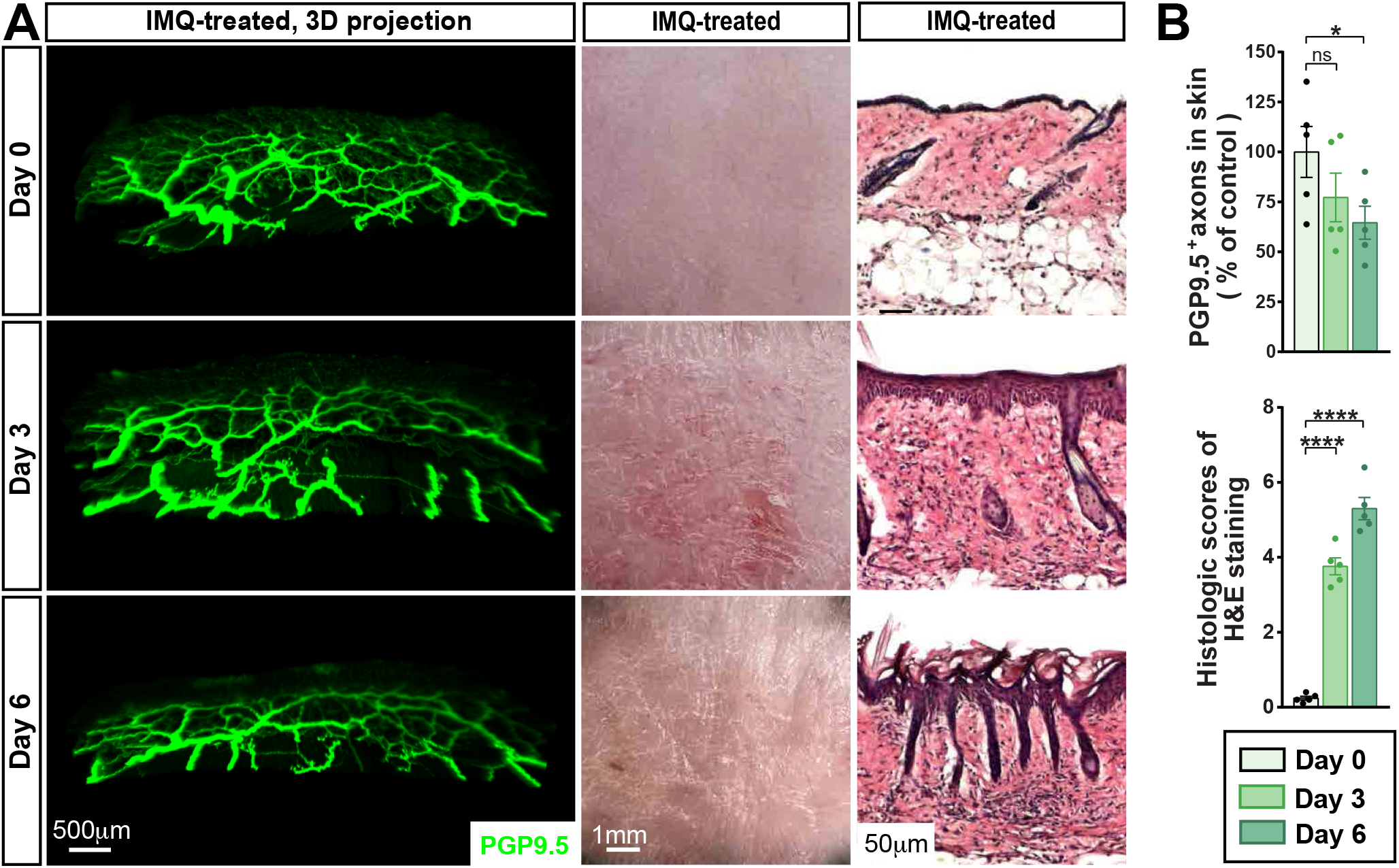
Assessment of skin inflammation induced by IMQ. Wild-type mice were subjected to daily and topical IMQ-treatment for six consecutive days. (**A**) Axonal projections throughout 1mm sections of skin were visualized whole mount immunolabeling with the pan-axonal marker PGP9.5 In parallel, the gross appearance of dorsal skin and corresponding H&E staining was recorded in untreated (Day 0) and following 3 or 6 days of daily treatment. (**B**) Quantification of axon number and pathology (by H&E). n = 5 mice, unpaired, two-tailed t-test.

**Figure S2.**
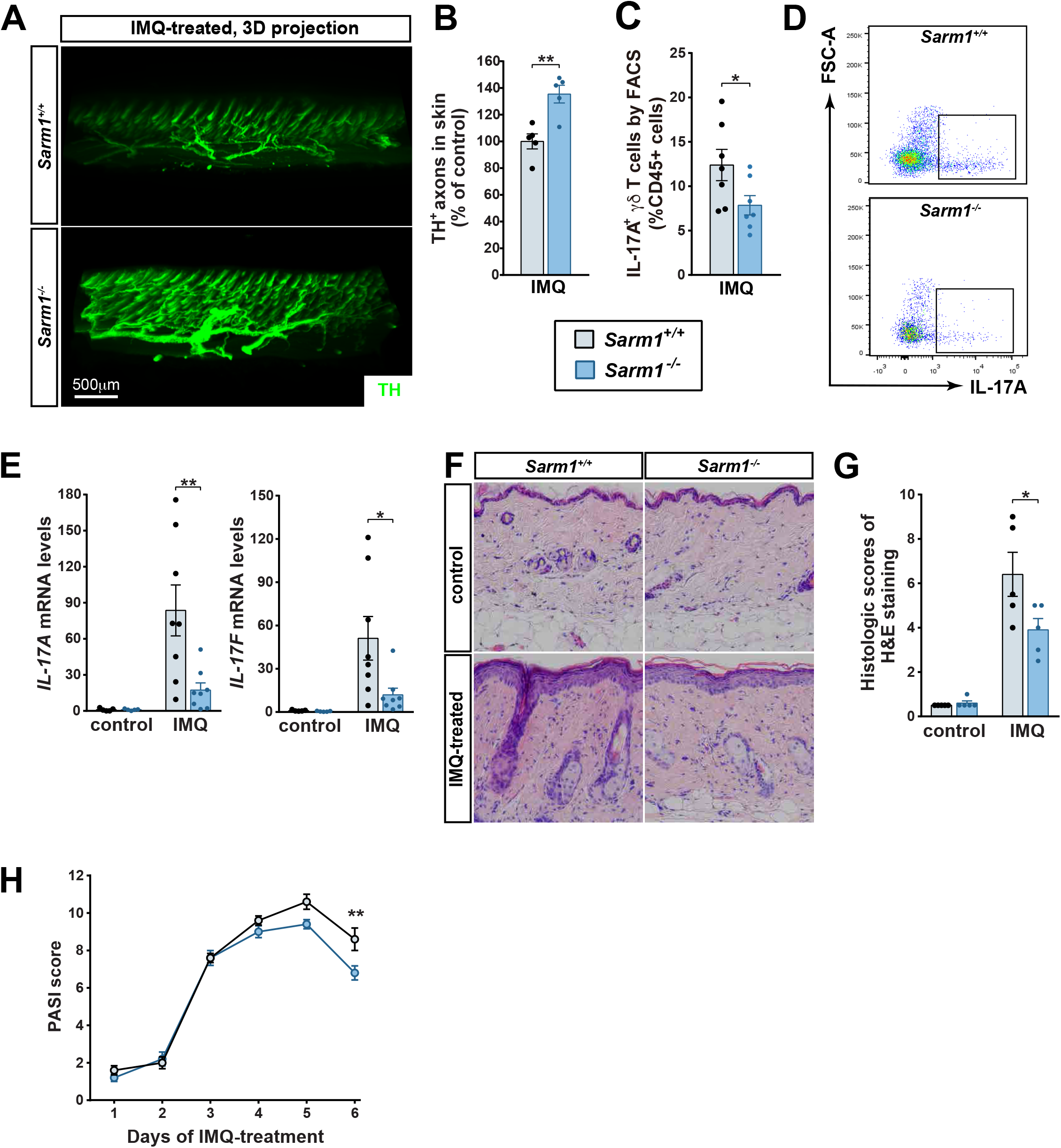
Blockade of axon degeneration in Sarm1 knockouts lessens IMQ-induced skin inflammation. (**A**) Mice of the indicated genotypes were subjected to daily IMQ-treatment for six consecutive days. Skin tissues were collected 24 hours after the last treatment of IMQ. Representative 3D-projection of skin treated as indicated and visualized by whole mount immunolabeling for TH. (**B**) TH-positive sympathetic axons were quantified. n = 5, mean ± SEM, unpaired, two-tailed t-test. (**C, D**) CD45^+^ CD3^+^ γδTCR^+^ IL-17A^+^ cells in skin were examined by FACS and quantified. *n* = 7 mice, mean ± SEM, unpaired, two-tailed t-test. (**E**) Expression levels of *IL-17A* and *IL-17F* in skin were determined by qPCR. *n* = 5 (control) or 8 (IMQ), means ± SEM, two-way ANOVA with Tukey’s post-hoc test. (**F**) Representative images of H&E-stained skin sections. (**G**) Histologic scores were determined. n = 5 mice, mean ± SEM, two-way ANOVA with Tukey’s post-hoc test. (**H**) PASI of mouse dorsal skin was evaluated every day before IMQ-treatment. *n* = 5 mice, mean ± SEM, two-way ANOVA with Sidak’s correction.

## Material and methods

### Mice

Experiments were performed on female mice between 8 and 10 weeks of age unless otherwise specified, and in accordance with approved protocols at Weill Cornell Medicine or Peking University. Wild type C57BL/6 mice were purchased from the Jackson Labs or Charles River International. Sarm1 knockout (Cat#JAX:018069, RRID:IMSR_JAX:018069) mice were purchased from Jackson Laboratory and backcrossed for one generation to wild type mice.

Heterozygous offspring were then intercrossed to generate Sarm1 WT and KO littermates. To generate *Th-Cre;Sarm1^fl/fl^* mice, *Th-Cre* (Cat#JAX:008601, RRID:IMSR_JAX:008601) mice were purchased from the Jackson Labs and crossed to a *Sarm1^fl/^fl* mouse that we previously generated (Sun *et al*., 2021). *Adrb2* wild type and knockout mice were generated by backcrossing the strain *Adrb1;Adrb2* (JAX 003810, RRID:IMSR_JAX:003810) to C57BL/6 for two generations to isolate heterozygous *Adrb2* mice for intercrosses to generate WT and KO mice.

#### Skin treatments

<u>Imiquimod</u>: Mice at 8 to 10 weeks of age received a daily topical dose of 5% Imiquimod (IMQ) cream (Aldara; 3M Pharmaceuticals) for 3 or 6 consecutive days, as indicated. On the day prior to the first treatment, mice were anesthetized and a 2cm^2^ region of hair on their backs was shaved with an electric razor followed and cleared of remaining hairs using a depilatory cream (Nair; Church & Dwight Co., Inc.). Subsequent IMQ treatments were performed under brief inhaled isoflurane anesthesia. <u>6-OHDA</u>: Each mouse was treated with 500μg 6-OHDA (dissolved in 200 μl sterile PBS containing 0.01% ascorbic acid; Sigma-Aldrich), administered by subcutaneous injection daily for 2 days. A 2cm^2^ patch of dorsal hair was removed under anesthesia 3 days after the last injection. Mice next received 5 daily treatments of IMQ.

#### Generation of Bone Marrow Chimeric Mice

For the establishment of bone-marrow chimeric mice (BMCM), wild-type recipient mice at 8 weeks of age received 11 grays of X-ray radiation across two equal doses, given 2 hours apart in the Peking University animal facility, according to published protocols (Eng et al., 2020). This treatment eliminated the bone marrow population of recipient mice. Bone marrow was harvested from donor mice of the indicated genotypes as previously published (Sun *et al*., 2021). In brief, femurs were harvested, flushed with RPMI-1640 media, followed by red blood cell lysis and filtration with a 75μm strainer. 2 × 10^6^ cells of the resultant bone marrow suspension were transplanted into each irradiated mouse, 30 minutes post-exposure, via intravenous injection. Chimeric mice were treated with IMQ 5 weeks after the transplantation.

### Whole mount immunolabeling and 3D Imaging

At the conclusion of each experiment dorsal skin was harvested following perfusion with 20 ml of phosphate-buffered saline (PBS)/heparin (10 μg/ml) and 20 ml of PBS/1% paraformaldehyde (PFA)/10% sucrose/heparin (10 μg/ml). Skin was removed and fixed in PBS/0.5%PFA at 4°C overnight. Next day, the tissue was washed with PBS at room temperature for 30 min., three times. All subsequent incubation steps were performed with gentle rotating.

Whole mount immunolabeling of axons in skin was performed using the iDISCO method (Renier et al., 2014). For detection of γδ T cells we used a modified to the iDISCO protocol where acetone replaced methanol for the initial tissue dehydration (Liu et al., 2020). Skin samples were washed sequentially with 20%, 40%, 60%, 80% and 100% methanol (in ddH_2_O) for 1 hour at room temperature. Samples were next washed in 100% methanal for 1 hour and then bleached with 5% H_2_O_2_ (1 volume of 30%H_2_O_2_ for 5 volumes of methanol, ice cold) at 4°C overnight. After bleaching, samples were re-hydrated sequentially in 80%, 60%, 40%, 20% methanol in ddH_2_O for 1 hour at room temperature. The tissues were then washed with PBS at room temperature for 30 minutes, twice, followed by PBS/0.2% Triton X-100/0.1% deoxycholate/10% DMSO, overnight. The tissues were blocked with PBS/0.2% Triton X-100/10% DMSO/5% normal donkey serum at room temperature overnight and then immunolabeled with intended primary antibodies (1:500 dilution) in PBS/0.1% Tween 20/heparin (10 μg/ml)/5% normal donkey serum at room temperature for 48 hours. The primary antibodies used in this study were rabbit anti-PGP9.5 (ProteinTech Group, RRID:AB_2210497), and rabbit anti-TH (Millipore, RRID:AB_390204).The tissues were then washed with PBS/0.1% Tween 20/heparin (10 μg/ml) at room temperature for 12 hours, with the fresh buffer added every 2 hours. The tissues were further immunolabeled with the corresponding Alexa Fluor dye–conjugated secondary antibodies (1:500 dilution; Thermo Fisher Scientific) in PBS/0.1% Tween 20/heparin (10 μg/ml)/5% normal donkey serum at room temperature for 48 hours. The tissues were washed with PBS/0.1% Tween 20/heparin (10 μg/ml) at room temperature for 24 hours, with fresh buffer added every 6 hours. The immunolabeled lung tissues were washed with PBS at room temperature for 2 hours and embedded in PBS/1% agarose before the optical clearing. The tissue blocks were incubated at room temperature with 20% methanol (diluted in ddH_2_O) for 1 hour three times, 40% methanol for 1 hour, 60% methanol for 1 hour, 80% methanol for 1 hour, 100% methanol for 1 hour, and 100% methanol overnight. The tissue blocks were then incubated at room temperature with the mixture of dichloromethane and methanol (2:1) for 3 hours, followed by 100% dichloromethane for 30 min three times. The tissue blocks were lastly incubated at room temperature with 100% dibenzyl ether for 12 hours twice.

For acetone-based iDISCO, skin tissues were treated at room temperature with 25% acetone (diluted in ddH_2_O) for 1 hour, 50% acetone for 3 hours, and 25% acetone for 1 hour. The tissues were washed with PBS at room temperature for 30 min twice, followed by PBS/30% sucrose for 2 hours. The tissues were bleached in PBS/30% sucrose/1%H_2_O_2_ at 4°C overnight. The tissues were then washed with PBS at room temperature for 30 min twice, followed by PBS/0.2% Triton X-100/0.1% deoxycholate/10% DMSO overnight. The tissues were blocked, incubated with primary and secondary antibodies, embedding, and cleared as described in iDISCO. The primary antibodies used were Armenian hamster anti-γδTCR (Biolegend, RRID:AB_313830) and rabbit anti-TH (Millipore, RRID:AB_390204).

### PASI scoring

Skin inflammation resulting from IMQ treatment was evaluated by three indexes: erythema, thickness, and scale, based on an established methodology (Wu et al., 2019). Briefly, each of these pathologies was scored from 0-4 (0: none, 1: mild, 2: moderate, 3: marked, 4: severe). The reported psoriasis area severity index (PASI) score is the sum of these three values.

### Histology

For histologic examination, skin was dissected after perfusion, postfixed with 4% PFA at 4°C overnight, and processed for hematoxylin and eosin (H&E) staining. Briefly, skin was embedded in paraffin, sectioned, stained with H&E to illustrate the nuclear and cytoplasmic architecture of each section. Scoring of IMQ-treated skin tissues was performed following the established Baker’s standard (Luo et al., 2016). Analysis was performed on 5 randomly selected regions (×20 magnification) from sections of each tissue.

### Quantification of mRNA expression

Expression of genes of interest was measured using Real-Time Quantitative PCR (qPCR). For analysis of skin, dorsal skin was acutely removed and homogenized in TRIzol (Thermo Fisher Scientific) by a rotor-stator homogenizer. Total RNA was subsequently isolated, reverse transcribed by PrimeScript RT reagent Kit (Takara) and quantified using SYBR Green Real-Time PCR Kit (Thermo Fisher Scientific). Gene-specific primer pairs are as follows: cyclophilin (F: TGGAGAGCACCAAGACAGACA, R: TGCCGGAGTCGACAATGAT), IL-17A (F: CAGGGAGAGCTTCATCTGTGT, R: GCTGAGCTTTGAGGGATGAT), IL-17F (F: CCCAGGAAGACATACTTAGAAGAAA, R: GCAAGTCCCAACATCAACAG).

### FACS Analysis

To quantify immune cell numbers in dorsal skin, a 1cm^2^ skin punch biopsy was performed, removed of subcutaneous adipose tissue, and incubated in RPMI-1640 (Sigma-Aldrich) containing Dispase ⊓ (Sigma-Aldrich), Golgi-Plug (1:1000, BD Biosciences), 2% FBS (Gibco), 1% HEPES (Corning, pH=7.2-7.6), 1%Penicillin / Streptavidin (Corning) at 37°C for 30 mins. Next, the dermis was separated from the epidermis and finely minced into small pieces with scissors, digested in RPMI-1640 with 1mg/mL Collagenase W (Sigma-Aldrich), Golgi Plug (1:1000), 0.5mg/mL Hyaluronidase (Sigma-Aldrich), 100ug/mL DNaseI (Sigma-Aldrich) at 37°C for 30 min and passed through a 70-μm cell strainer. This resulted in a single cell suspension. Cells were then stimulated with 75ng/mL phorbol 12-myristate 13-acetate/ionomycin (Santa Cruz) for 4 hours in the presence of GolgiPlug (1:1000), then pre-incubated with Fc Block (eBioscience) and stained with surface markers (CD45,CD3, γδ TCR) and intracellular marker IL17A successively. Samples were analyzed using a BD LSRFortessa (BD Biosciences). FACS data were analyzed by FlowJo (https://www.flowjo.com).

### In Vitro Treatment

Skin subcutaneous lymph nodes were homogenized manually through 70-μm cell strainers. Cells were stained with antibodies to CD45 (Biolegend, RRID: AB_312979), CD3 (Biolegend, RRID:AB_312663) and γδ TCR (Biolegend, RRID:AB_313830), and sorted using a BD FACSAria (BD Biosciences). Primary γδT cells [CD45+CD3+γδ TCR+] were suspended in RPMI-1640 containing 10% heat-inactivated fetal bovine serum/penicillin (100 U/ml)/ streptomycin (100 μg/ml) and incubated at 37°C for 2hrs. Cells were subsequently treated with 20 μmol/L of either norepinephrine (Sigma), Formoterol (Sigma), or Clenbuterol (Sigma). Total RNA was then extracted using the RNeasy Mini Kit (Qiagen) and analyzed by qPCR as above.

## Notes

### Competing Interest Statement

The authors have declared no competing interest.

## References

Chiu, I.M., von Hehn, C.A., and Woolf, C.J. (2012). Neurogenic inflammation and the peripheral nervous system in host defense and immunopathology. Nat Neurosci 15, 1063–1067. 10.1038/nn.3144.

Croese, T., Castellani, G., and Schwartz, M. (2021). Immune cell compartmentalization for brain surveillance and protection. Nat Immunol 22, 1083–1092. 10.1038/s41590-021-00994-2.

Eng, J., Orf, J., Perez, K., Sawant, D., and DeVoss, J. (2020). Generation of bone marrow chimeras using X-ray irradiation: comparison to cesium irradiation and use in immunotherapy. J Biol Methods 7, e125. 10.14440/jbm.2020.314.

Essuman, K., Summers, D.W., Sasaki, Y., Mao, X., DiAntonio, A., and Milbrandt, J. (2017). The SARM1 Toll/Interleukin-1 Receptor Domain Possesses Intrinsic NAD(+) Cleavage Activity that Promotes Pathological Axonal Degeneration. Neuron 93, 1334–1343 e1335. 10.1016/j.neuron.2017.02.022.

Flutter, B., and Nestle, F.O. (2013). TLRs to cytokines: mechanistic insights from the imiquimod mouse model of psoriasis. Eur J Immunol 43, 3138–3146. 10.1002/eji.201343801.

Gaskill, P.J., and Khoshbouei, H. (2022). Dopamine and norepinephrine are embracing their immune side and so should we. Curr Opin Neurobiol 77, 102626. 10.1016/j.conb.2022.102626.

Gold, M.H., Holy, A.K., and Roenigk, H.H., Jr. (1988). Beta-blocking drugs and psoriasis. A review of cutaneous side effects and retrospective analysis of their effects on psoriasis. J Am Acad Dermatol 19, 837–841.

Ingolia, N.T., and Murray, A.W. (2007). Positive-feedback loops as a flexible biological module. Curr Biol 17, 668–677. 10.1016/j.cub.2007.03.016.

Itohara, S., Mombaerts, P., Lafaille, J., Iacomini, J., Nelson, A., Clarke, A.R., Hooper, M.L., Farr, A., and Tonegawa, S. (1993). T cell receptor delta gene mutant mice: independent generation of alpha beta T cells and programmed rearrangements of gamma delta TCR genes. Cell 72, 337–348. 10.1016/0092-8674(93)90112-4.

Jin, L., Zhang, J., Hua, X., Xu, X., Li, J., Wang, J., Wang, M., Liu, H., Qiu, H., Chen, M., et al. (2022). Astrocytic SARM1 promotes neuroinflammation and axonal demyelination in experimental autoimmune encephalomyelitis through inhibiting GDNF signaling. Cell Death Dis 13, 759. 10.1038/s41419-022-05202-z.

Kagi, D., Ledermann, B., Burki, K., Seiler, P., Odermatt, B., Olsen, K.J., Podack, E.R., Zinkernagel, R.M., and Hengartner, H. (1994). Cytotoxicity mediated by T cells and natural killer cells is greatly impaired in perforin-deficient mice. Nature 369, 31–37. 10.1038/369031a0.

Klein Wolterink, R.G.J., Wu, G.S., Chiu, I.M., and Veiga-Fernandes, H. (2022). Neuroimmune Interactions in Peripheral Organs. Annu Rev Neurosci 45, 339–360. 10.1146/annurev-neuro-111020-105359.

Ko, K.W., Milbrandt, J., and DiAntonio, A. (2020). SARM1 acts downstream of neuroinflammatory and necroptotic signaling to induce axon degeneration. J Cell Biol 219. 10.1083/jcb.201912047.

Liu, T., Yang, L., Han, X., Ding, X., Li, J., and Yang, J. (2020). Local sympathetic innervations modulate the lung innate immune responses. Sci Adv 6, eaay1497. 10.1126/sciadv.aay1497.

Luo, D.Q., Wu, H.H., Zhao, Y.K., Liu, J.H., and Wang, F. (2016). Original Research: Different imiquimod creams resulting in differential effects for imiquimod-induced psoriatic mouse models. Exp Biol Med (Maywood) 241, 1733–1738. 10.1177/1535370216647183.

Martini, R., and Willison, H. (2016). Neuroinflammation in the peripheral nerve: Cause, modulator, or bystander in peripheral neuropathies? Glia 64, 475–486. 10.1002/glia.22899.

Mombaerts, P., Clarke, A.R., Hooper, M.L., and Tonegawa, S. (1991). Creation of a large genomic deletion at the T-cell antigen receptor beta-subunit locus in mouse embryonic stem cells by gene targeting. Proc Natl Acad Sci U S A 88, 3084–3087. 10.1073/pnas.88.8.3084.

Panneerselvam, P., Singh, L.P., Selvarajan, V., Chng, W.J., Ng, S.B., Tan, N.S., Ho, B., Chen, J., and Ding, J.L. (2013). T-cell death following immune activation is mediated by mitochondria-localized SARM. Cell Death Differ 20, 478–489. 10.1038/cdd.2012.144.

Pasparakis, M., Haase, I., and Nestle, F.O. (2014). Mechanisms regulating skin immunity and inflammation. Nat Rev Immunol 14, 289–301. 10.1038/nri3646.

Ren, K., and Dubner, R. (2010). Interactions between the immune and nervous systems in pain. Nat Med 16, 1267–1276. 10.1038/nm.2234.

Renier, N., Wu, Z., Simon, D.J., Yang, J., Ariel, P., and Tessier-Lavigne, M. (2014). iDISCO: a simple, rapid method to immunolabel large tissue samples for volume imaging. Cell 159, 896–910. 10.1016/j.cell.2014.10.010.

Roosterman, D., Goerge, T., Schneider, S.W., Bunnett, N.W., and Steinhoff, M. (2006). Neuronal control of skin function: the skin as a neuroimmunoendocrine organ. Physiol Rev 86, 1309–1379. 10.1152/physrev.00026.2005.

Russell, J.H., and Ley, T.J. (2002). Lymphocyte-mediated cytotoxicity. Annu Rev Immunol 20, 323–370. 10.1146/annurev.immunol.20.100201.131730.

Sambashivan, S., and Freeman, M.R. (2021). SARM1 signaling mechanisms in the injured nervous system. Curr Opin Neurobiol 69, 247–255. 10.1016/j.conb.2021.05.004.

Sun, Y., Wang, Q., Wang, Y., Ren, W., Cao, Y., Li, J., Zhou, X., Fu, W., and Yang, J. (2021). Sarm1-mediated neurodegeneration within the enteric nervous system protects against local inflammation of the colon. Protein Cell 12, 621–638. 10.1007/s13238-021-00835-w.

van der Fits, L., Mourits, S., Voerman, J.S., Kant, M., Boon, L., Laman, J.D., Cornelissen, F., Mus, A.M., Florencia, E., Prens, E.P., and Lubberts, E. (2009). Imiquimod-induced psoriasis-like skin inflammation in mice is mediated via the IL-23/IL-17 axis. J Immunol 182, 5836–5845. 10.4049/jimmunol.0802999.

Voskoboinik, I., Whisstock, J.C., and Trapani, J.A. (2015). Perforin and granzymes: function, dysfunction and human pathology. Nat Rev Immunol 15, 388–400. 10.1038/nri3839.

Wang, Q., Zhang, S., Liu, T., Wang, H., Liu, K., Wang, Q., and Zeng, W. (2018). Sarm1/Myd88-5 Regulates Neuronal Intrinsic Immune Response to Traumatic Axonal Injuries. Cell Rep 23, 716–724. 10.1016/j.celrep.2018.03.071.

Wlaschin, J.J., Gluski, J.M., Nguyen, E., Silberberg, H., Thompson, J.H., Chesler, A.T., and Le Pichon, C.E. (2018). Dual leucine zipper kinase is required for mechanical allodynia and microgliosis after nerve injury. Elife 7. 10.7554/eLife.33910.

Wu, D.H., Zhang, M.M., Li, N., Li, X., Cai, Q.W., Yu, W.L., Liu, L.P., Zhu, W., and Lu, C.J. (2019). PSORI-CM02 alleviates IMQ-induced mouse dermatitis via differentially regulating pro-and anti-inflammatory cytokines targeting of Th2 specific transcript factor GATA3. Biomed Pharmacother 110, 265–274. 10.1016/j.biopha.2018.11.092.

Yatim, N., Cullen, S., and Albert, M.L. (2017). Dying cells actively regulate adaptive immune responses. Nat Rev Immunol 17, 262–275. 10.1038/nri.2017.9.

